# A re-interpretation of the experimental data of Shinder and Taube “Three-dimensional Tuning of Head Direction Cells in Rats”, Journal of Neurophysiology 121(1), 2019

**DOI:** 10.1101/559336

**Authors:** Jean Laurens, Dora E. Angelaki

## Abstract

In a recent study, Shinder and Taube (2019) concluded that head direction cells in the anterior thalamus of rats are tuned to one-dimensional (1D, yaw-only) motion exclusively, in contrast to recent findings in bats (Finkelstein et al. 2015), mice (Angelaki et al. 2016; Cham et al. 2017; Laurens et al. 2017), and rats (Page et al. 2017). Here we re-interpret the author’s experimental results using model comparison and demonstrate that, contrary to their conclusions, their data actually supports the dual-axis rule (Page et al. 2017) and tilted azimuth model (Laurens and Angelaki 2018), where head direction cells use gravity to integrate 3D rotation signals about all cardinal axes of the head. We further show that this study is inconclusive regarding the presence of vertical orientation tuning; i.e. whether head direction cells encode 3D orientation in the horizontal and vertical planes conjunctively. Using model simulations, we demonstrate that, even if 3D tuning existed, the experimental protocol and data analyses used by Shinder and Taube (2019) would not have revealed it. We conclude that the actual experimental data of Shinder and Taube (2019) are compatible with the 3D properties of head direction cells discovered by other groups, yet incorrect conclusions were reached because of incomplete and qualitative analyses.

## Introduction

Head direction cells (HDC) track allocentric head orientation, similar to a ‘neuronal compass’. Since their discovery (Taube et al. 1990a,b), most studies have focused on how HDC encode head orientation in the horizontal plane (azimuth) when animals explore horizontal surfaces. Initial studies during 2D motion concluded that HDC treat vertical walls as an extension of the floor (**Fig. 1A**) and postulated that the HDC system tracks head orientation by integrating yaw rotations (i.e. left/right rotations in the egocentric head horizontal plane) exclusively (‘Yaw-only’, YO, model; **Fig. 1A**; Stackman et al. 2000, Calton and Taube 2005). Furthermore, these studies observed that HDC lose their tuning when animals walk in an inverted orientation, e.g. on the ceiling in **Fig. 1A**.

**Figure 1:**
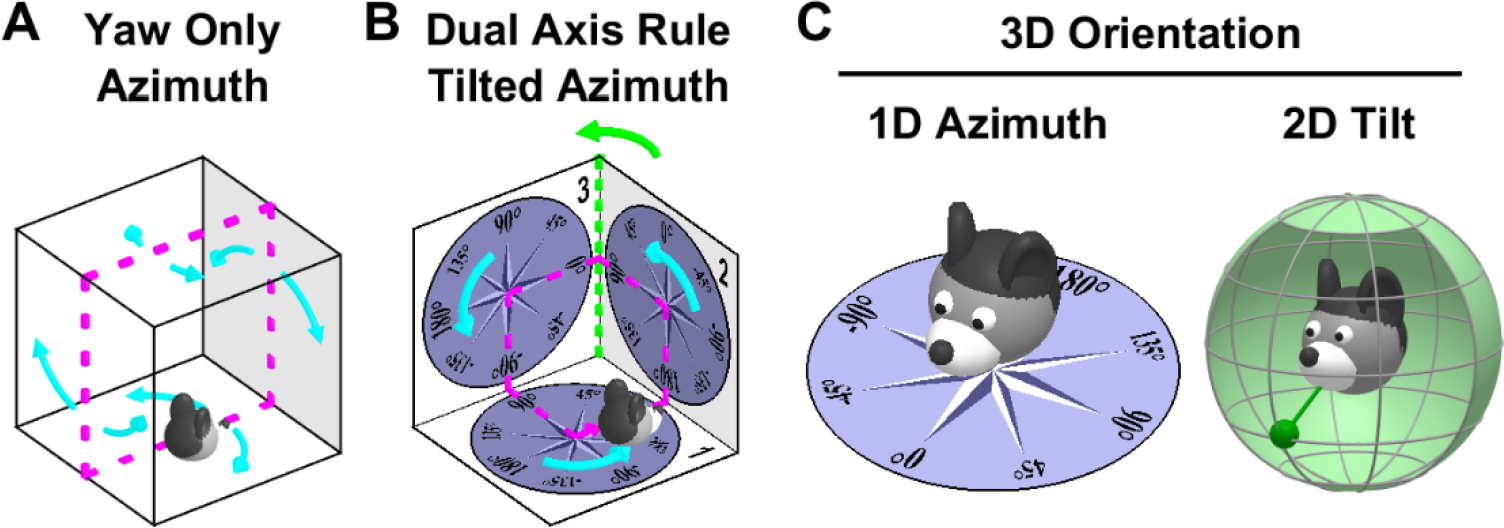
Fundamental difference between the Yaw-only (YO) and tilted azimuth (TA) models. **A:** The Yaw-only (YO) azimuth model (Calton and Taube 2005, Shinder and Taube 2019) assumes that vertical surfaces are treated as extension of the floor. Azimuth is computed by integrating yaw (i.e. left/right, cyan) rotations only, irrespective of the allocentric orientation of the locomotion surfaces. This model is sufficient to track head orientation when walking on the floor, ceiling and opposite walls of a cage (magenta). **B:** The difference between the YO and tilted azimuth (TA; Laurens and Angelaki 2018)/dual axis rule (Page et al. 2017) model arises when movement is not restricted to 2D. In this example, the animal completes a circular trajectory (magenta) that includes three left-hand yaw turns (cyan). The YO model cannot maintain allocentric consistency, as it will only register a total azimuth rotation angle of 270°, in contradiction with the fact that the animal has returned to its initial orientation. The ‘missing’ rotation not registered by the YO model occurs when the animal transitions from surface 2 to 3 (green). This is the 2^nd^ axis of the dual axis rule: Tracking 3D orientation correctly requires updating azimuth <u>also</u> when the locomotion surface (or the head horizontal plane) crosses vertical surfaces, i.e., during rotations about an earth-vertical axis. The equivalent TA model (Laurens and Angelaki 2018, see **Fig. 3A** for definition) assumes that azimuth is measured in the compasses drawn on the three surfaces and yields a correct total rotation of 360°. Thus, the YO model of Shinder and Taube (2019) and the TA/dual axis model of Page et al. (2017) and Laurens and Angelaki (2018) are consistent with each other <u>only</u> when movement is restricted to 2D (e.g., magenta trajectory in A). The YO model would fail to track azimuth during 3D movements (as shown in B; see also Laurens and Angelaki, 2018). Thus, by supporting the YO model, Shinder and Taube (2019) imply that the HD system will fail during 3D movements on the surfaces of a cube, which is inconsistent with experimental results (Page et al. 2017). **C:** Decomposition of 3D orientation into 1D tilted azimuth and 2D tilt, as proposed by (Laurens et al. 2016, Laurens and Angelaki 2018). Azimuth measures 1D orientation relative to allocentric horizontal landmarks and has a circular topology. Tilt measures 2D orientation of the head relative to gravity, or equivalently the egocentric orientation of the gravity vector (left panel: green pendulum) and has a spherical topology. This model generalizes Finkelstein's toroidal (Yaw-Pitch) model (Finkelstein et al. 2015) to 3D.

Recent studies by several other groups, however, have shed additional insight about the 3D properties of HDC. Experimental (Page et al. 2017) and theoretical (Jeffery et al. 2013; Laurens and Angelaki 2018) studies have shown that HDC track azimuth by combining egocentric rotations about the head’s three cardinal axes, not only yaw, with gravity signals. This process, which has been called ‘dual axis rule’ in (Page et al. 2017, **Fig. 1B**) and tilted azimuth (TA) model in (Laurens and Angelaki 2018), was confirmed experimentally by HDC recordings in the anterior thalamus of rats (Page et al. 2017). Furthermore, a series of studies have shown that HDC in bats (Finkelstein et al. 2016), mice (Angelaki et al. 2016; Cham et al. 2017; Laurens et al. 2017) and likely macaques (Laurens et al. 2016) can also represent head tilt, i.e. head orientation in vertical planes. By encoding 1D azimuth and 2D tilt conjunctively, the HDC system can thus represent 3D orientation (**Fig. 1C**).

Yet, a recent study of HDC in rats (Shinder and Taube 2019) has challenged the recent evidence for a 3D organization of the HDC system by resurrecting the YO model. Based on a series of 3D passive rotation protocols, the authors have re-iterated the conclusions from previous studies, i.e. that HDC: (1) are updated by yaw rotations only; and (2) don’t exhibit any tilt (vertical orientation) tuning. These conclusions were made without quantitative analyses and model comparison, and without discussing the contradiction to recent findings from other groups.

Why do Shinder and Taube’s conclusions differ from recent findings in an array of mammalian species (Page et al. 2017, Laurens et al. 2016, Angelaki et al. 2016; Cham et al. 2017; Laurens et al. 2017; Finkelstein et al. 2015)? We have re-examined their data quantitatively using comparisons with model simulations, and reached entirely different conclusions. Specifically, we find that (1) the HDC responses of Shinder and Taube (2019) cannot be predicted by the YO model, in direct conflict with their conclusion, but are in fact supportive of the dual axis/TA azimuth model. Importantly, the YO and the dual axis/TA models are mutually exclusive. The dual axis rule, by definition, implies that 3D, not just yaw, rotation signals processed by gravity are necessary to update and maintain the azimuth tuning of HDC.

Regarding their second conclusion, we demonstrate that (2) the experimental protocol and data analyses used by Shinder and Taube (2019) entangle azimuth and putative tilt tuning in a way that would conceal the existence of tilt tuning; thus making their study essentially inconclusive in this regard.

Here we present this analysis, first focusing on azimuth coding by providing a quantitative comparison between predictions of the two azimuth (TA and YO) models (which Shinder and Taube never did). We next present model simulations demonstrating that, even if tilt tuning existed, it would not have been obvious using the authors’ experimental protocol and data analyses. We start with a brief description of the two azimuth tuning models.

### Shinder and Taube: HDC are tuned to azimuth only based on a YO model that leads to a hemitorus/ellipsoidal tuning scheme

According to Shinder and Taube (2019; see also Stackman and Taube, 2000), HDC are tuned to azimuth only, as follows: (1) azimuth is computed by always integrating exclusively yaw (head-fixed left-right) rotations (Fig. **1A**, cyan; referred to as the ‘YO model’); (2) on sloped surfaces, a HD cell fires as if the surfaces were extensions of the floor; (3) azimuth tuning is lost at tilt angles larger than 90°.

To conceptualize these properties, Shinder and Taube (2019) refer to a “hemitorus” (which, should be noted, is unrelated to Finkelstein’s (2015) toroidal model) and “ellipsoid”, illustrated graphically in **Fig. 2**. The hemitorus is produced by plotting the cell’s tuning curve in a polar plot on the head’s yaw plane (**Fig. 2A**, dark blue: response on the earth-horizontal and earth-vertical planes) as the head tilts ±90° about an axis perpendicular to the PD (**Fig. 2A**, red). Note that the hemitorus restricts head tilt to 90°, such as to imply the loss of tuning when the head is inverted (allowing the head’s horizontal plane to rotate fully (±180°) would form a full torus). The ellipsoidal model follows a similar rationale (**Fig. 2B**), now describing tilts about an axis parallel to the PD (red line).

**Figure 2:**
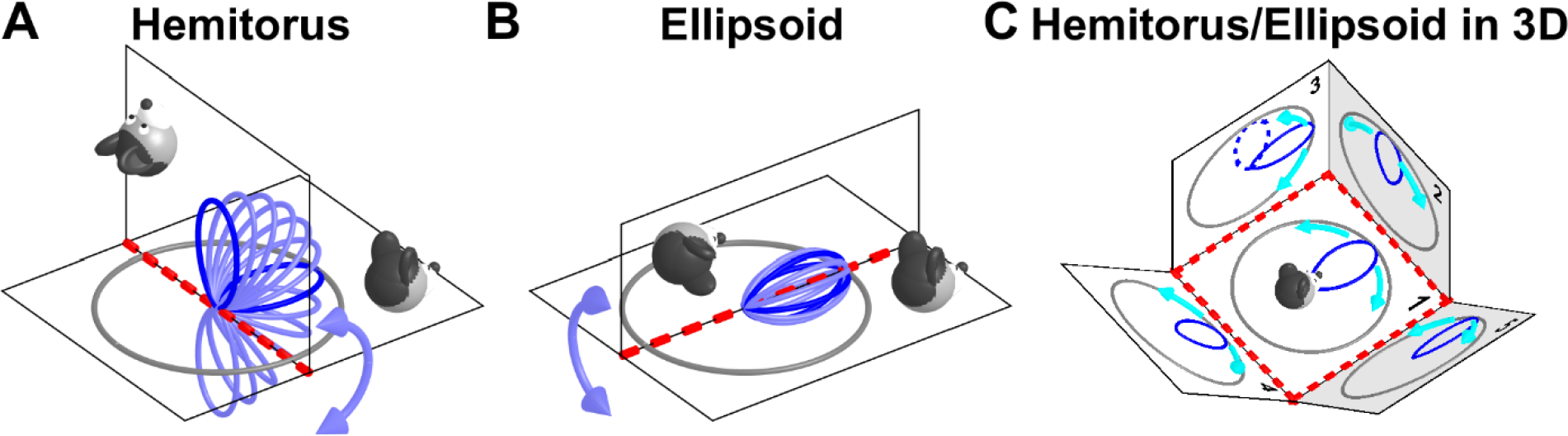
Hemitorus and ellipsoid schematics corresponding to the YO model. **A**: HDC tuning curves at different head orientations between nose-up and nose-down (light blue) form a hemitorus when the tilt axis (red) is perpendicular to the cell’s PD. Dark blue: tuning curves when animal oriented as in the drawing. **B**: HDC tuning curves form an ellipsoid when the tilt axis (red) is now parallel to the cell’s PD. The orientations shown in A and B do not cover the full 3D space, therefore the YO model is incomplete and inadequate to maintain 3D orientation consistently, as shown in C. **C**: Predicted tuning curves on multiple planes spanning 3D space. The plane marked ‘1’ is earth-horizontal. The tuning curves shown on earth-vertical planes ‘2’ and ‘3’ are predicted by the hemitorus and ellipsoid YO model, respectively. The tuning curves on planes ‘4’ and ‘5’, which are tilted by 45°, are also predicted by same YO model (where azimuth is updated by yaw rotations only, cyan arrow) during transitions from plane 1. In fact, the YO model allows tracking azimuth accurately when the head stays in one plane or transitions between plane 1 and any other plane (a ‘null’ rotation; see **Fig. 3A**). However, the YO model fails whenever the animal’s movement is such that updating azimuth requires use of the 2^nd^ updating rule (rotation about an earth-vertical axis). The broken line on plane 3 shows the tuning curve that would result from transitioning from panel 2 to panel 3 based on the YO model. Note that the same physical orientation on plane 3 would be registered differently by the YO model depending on whether the animal originated from plane 1 or plane 2. Thus, the YO model would not maintain allocentric invariance when animals move in 3D.

Note that the hemitorus and ellipsoidal models are not only qualitative in nature, but also incomplete; they do not consider truly 3D movements: each covers a 2D range of movements (the hemitorous model deals with movements that can be expressed as pitch followed by yaw and the ellipsoid model deals with roll + yaw), and together the two models cover only two 2D subspaces of the ensemble of 3D head rotations, leaving out most 3D orientations. Most importantly, the YO model of Shinder and Taube (2019) would completely loose allocentric invariance for any movement that would bring the animal’s head outside these two planes (**Fig. 1B**; **Fig. 2C**; see Laurens and Angelaki 2018 for details).

This problem is illustrated by plotting the predicted firing rate of a HDC when the animal walks on a variety of planes (**Fig. 2C**). In this figure, the hemitorous is used to predict the tuning curves when the animal transitions between surfaces 1,2 and 4 and the ellipsoid is used to predict the tuning curves when the animal transitions between surfaces 1,3 and 5. What would happen if the animal transitioned between other pairs of surfaces? E.g., if the animal transitioned from surface 2 (where the tuning curve peaks when the animal faces ‘upward’) to surface 3, the YO model would not update; thus, the resulting tuning curve (**Fig. 2C**, broken blue line) would still point ‘upward’, in contradiction with the prediction of the ellipsoid model (solid line). This is because, according to Shinder and Taube’s YO model, azimuth would not get updated during this transition from surface 2 to surface 3. Here lies exactly the problem with the YO model: it cannot handle 3D rotations and is not inherently consistent when movements are not restricted to rotations about axes parallel and perpendicular to HDC PDs.

### Model of HDC during 3D motion: the TA/dual axis rule

In contrast to the YO model, updating TA azimuth must follow the dual axis rule (Page et al. 2017; **Fig. 3A**), by integrating <u>both</u> yaw (**Fig. 3A**, cyan) <u>and</u> earth-vertical axis (**Fig. 3A**, green) rotations. One can think of any arbitrary rotation being decomposed into a frame formed by the head-fixed yaw axis, the earth-vertical axis and the null axis (**Fig. 3A**, red). Both yaw and earth-vertical axis rotations should update TA (in contrast to the YO model, which is updated only during yaw rotations); only rotations about the null axis do not update TA. By definition, the null axis is orthogonal to the yaw and earth-vertical axes (which are themselves not necessarily orthogonal) and therefore lies at the intersection of the head-horizontal and earth-horizontal planes. Thus, the TA at any 3D head orientation can be computed by placing a compass in the head-horizontal plane, and orienting it such that it matches the earth-horizontal compass at the level of the null axis (**Fig. 3A**, red; e.g., both compasses read ±90° along this line). Further details about the TA and dual axis rule models can be found in (Page et al. 2017, Laurens and Angelaki 2018).

**Figure 3:**
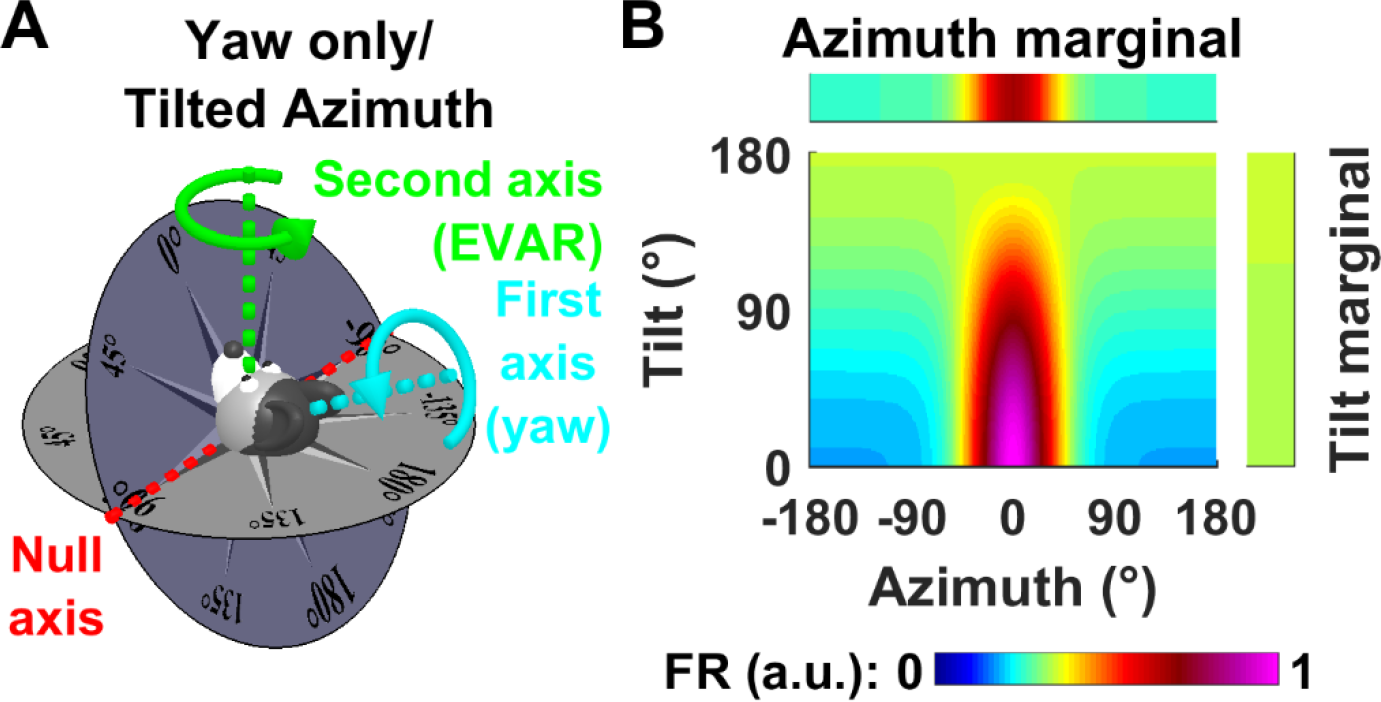
Tilted azimuth (TA)/dual axis rule model and dependence of azimuth tuning on tilt angle. **A:** Definition of the TA model: Azimuth is measured in a compass affixed to the head-horizontal plane (dark blue), oriented so that it matches the earth-horizontal compass (grey) along the null axis (red line). In this model, azimuth is updated <u>both</u> during head-fixed yaw rotations (cyan arrow) and by rotations around an earth-vertical axis (green). This 2^nd^ updating rule is necessary for the TA compass to maintain allocentric invariance (i.e., consistency with the earth-horizontal compass). Rotations about the null axis (red) don’t affect TA. Note that <u>the TA and YO models are entirely distinct when considering 3D orientation</u>, because the latter does not include the dual axis rule and only updates during yaw rotations. **B:** Model tilt-dependent azimuth tuning curve, with a PD at 0° azimuth (magenta), illustrating tilt-dependent azimuth tuning strength. The marginal distributions indicate the average firing across all tilt angles (azimuth marginal) and across all azimuth (tilt marginal; which is uniform). Note that a tilt angle-dependent azimuth tuning results in an un-tuned tilt marginal (i.e., the model cell is not tuned to tilt).

Of course, when the head is upright, yaw and earth-vertical axes are identical. When the head is upside-down, these two axes are opposite. Thus, TA is not defined in this position, accounting perhaps for the fact that the cell isn’t tuned to azimuth when upside-down.

### Tilt-dependent azimuth tuning

Next, for both the YO and TA models, we add a property, which for now can be considered an assumption (but we know from our own experimental data to be true; Laurens et al. 2017): the magnitude of azimuth tuning is largest in upright orientation and decreases as a function of head tilt. We illustrate this assumption by representing the firing rate of a model cell as a function of azimuth (**Fig. 3B**, abscissa) and tilt (**Fig. 3B**, ordinate) as a color map. On average across all tilt angles (upper marginal distribution), the cell is tuned to azimuth. Note, however, that the average firing rate as a function of tilt across all azimuths (right marginal distribution) is uniform. As will be shown later, we define tilt tuning as the average firing rate across all azimuths and therefore, by such a definition, a cell’s azimuth tuning can depend on tilt angle, without the cell being tuned to tilt.

Mathematically, we model azimuth tuning curve when the head is upright by a von Mises function. For simplicity, we set the cell’s preferred azimuth to 0° and its peak firing rate to 1:

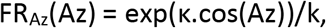

where Az is the azimuth angle, κ is the von Mises function’s parameter and k is a normalizing constant set such that FR_Az_(0) = 1. The hypothesized tilt-dependent modulation of the tuning strength can be parametrized as follows (α: tilt angle):

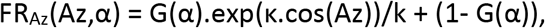

where G(α) = k_G_.(1-sin^2^(α/2)) is a tilt-dependent gain function with parameter k_G_ and k is the average of exp(κ.cos(Az)) across all values of Az, such that the average of FR_Az_(Az,α) across all Az is equal to 1 independently of α. Note that, in upside-down position (α=180°), G(α) is equal to zero and the cell’s firing rate is set to 1. Thus, even though TA is not defined in upside-down position, the cell’s firing rate can still be computed continuously.

### YO vs. TA models

The difference between the dual-axis rule and the YO model is also illustrated **Fig. 2C**. The dual axis rule predicts that azimuth will be updated by yaw rotations when the animal remains in a given surface, similar to the YO model. Furthermore, like the YO model, the dual axis rule predicts that azimuth should not be updated when transitioning between surface 1 and any other surface, since these transitions correspond to the null axis (marked as red in **Fig. 2C**). However, the dual-axis rule predicts that azimuth should be updated by the second rule when transitioning between vertical surfaces, e.g. 2 and 3 in **Fig. 2C**, which would allow maintaining consistency between the tuning curves shown on all surfaces.

In their publication, Shinder and Taube (2019) do not acknowledge that the YO and TA model are inconsistent with each other. In fact, they repetitively suggest that the two are the same, which is absolutely incorrect. Specifically:

- The YO and TA models have in common that azimuth is measured on a compass affixed to the head-horizontal plane. In fact, the two models are equivalent for a restricted range of 2D movements (i.e. those in **Fig. 2A,B** with less than 90° tilt). However, because the YO model lacks the second axis rule, it predicts that HD responses will lose consistency during 3D rotations. The TA model can handle any 3D re-orientation, but the YO model fails saliently.
- The YO/hemitorus/ellipsoidal model incorporates the added assumption that HD tuning vanishes abruptly when tilt angle exceeds 90°. The TA model is undefined in upside-down orientation (i.e. at a tilt angle of 180°) but otherwise doesn’t make any assumption about the range in which HDC encode azimuth. We added a tilt-dependent gain modulation in the TA model, whereby HDC responses decrease continuously when head tilts.

Replacing that azimuth tuning vanishes when animals are inverted with a cutoff at a tilt angle of 90°, with a smooth and continuous dependence, is but a small detail. To be able to directly compare the other more important predicted differences of the YO versus TA model, we assumed the same smooth tilt angle dependence for both models. Thus, all simulations shown here differ only on the fundamental difference between YO and TA models: the 2^nd^ dual axis updating rule.

### Simulations of TA and YO models: HDC responses support the dual axis/TA model

In order to determine whether Shinder and Taube’s data support the dual axis/TA rule or the yaw-only model, we simulated the responses of model HDC encoding either YO or TA azimuth using the whole range of manipulations performed in (Shinder and Taube 2019). These simulations were compared to the average responses reported by the authors.

To determine each model’s parameter κ and k_G_, we fitted them to the average HD response reported in all manipulations. We found that the best fitting parameters were similar in both models: κ = 4.6 and k_G_ = 0.46 for the TA model and κ = 6.3 and k_G_ = 0.42 for the YO model.

#### HDC integrate 3D rotation signals, and not only yaw

In the first manipulation (**Fig. 4A**), the head is upright and the rotation axis coincides with the head’s yaw axis. Therefore, the trajectories are the same in both YO and TA models (‘Trajectory’ panel, TA: blue; Yaw-only: cyan) and the predicted tuning curves (‘Tuning Curves’ panel) are similar (although the parameters of both models differ slightly, this difference is negligible in practice). Both predictions are highly correlated (TA: ρ=0.99; YO: ρ=0.97) with the average response measured across cells (‘Tuning Curves’ panel, black).

**Figure 4:**
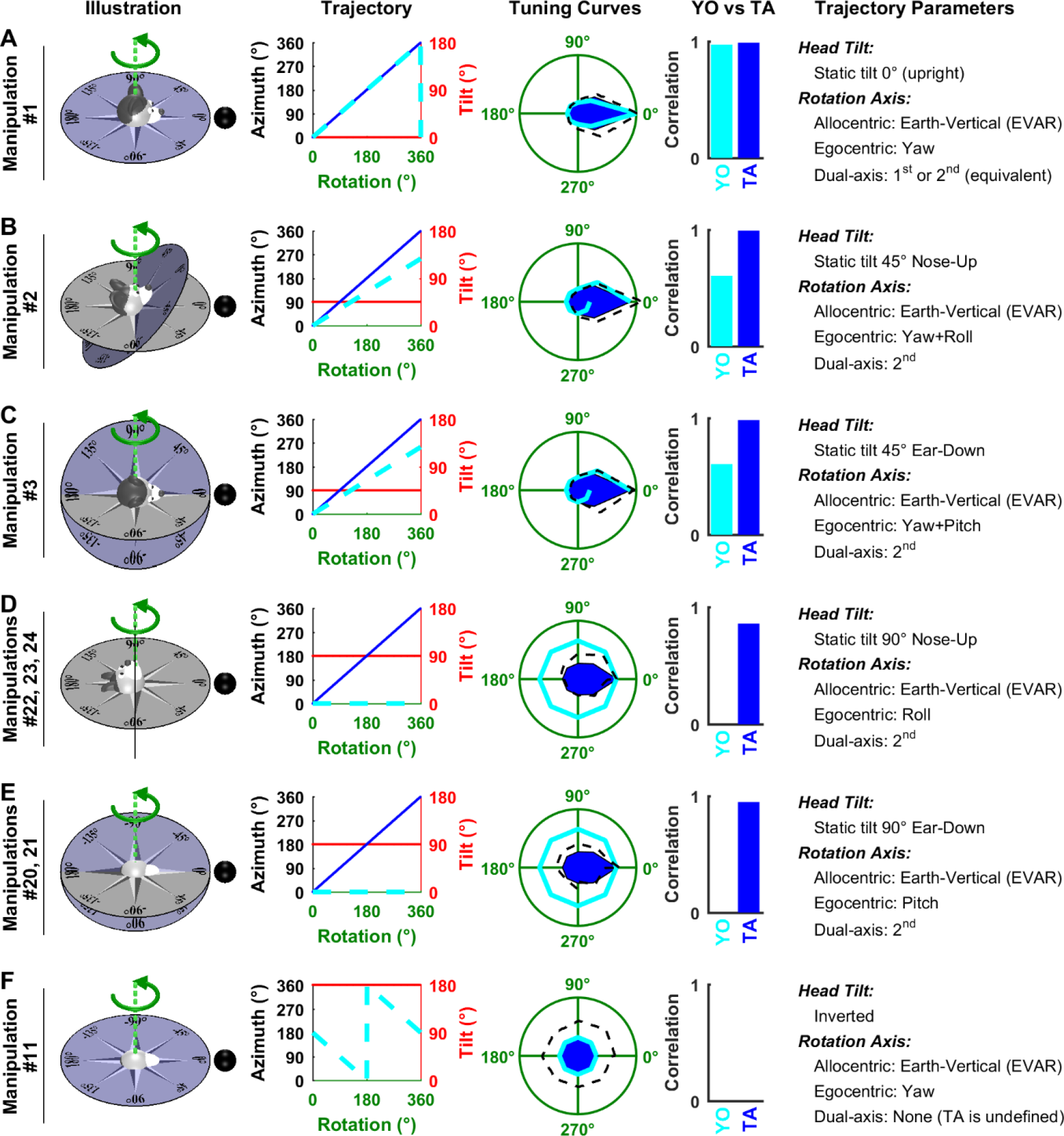
Analysis of Shinder and Taube’s (2019) manipulations: rotations around an earth-vertical axis. Experimental conditions in (Shinder and Taube 2019) are named ‘manipulations’ here, numbered 1 to 27 (see their Fig. 3). Each line corresponds to one manipulation, or to multiple manipulations that follow the same trajectory (with different starting positions). **‘Illustration’:** representation of the motion trajectories with the head at 0°. The grey and blue azimuth frames represent an earth-fixed and TA compass, respectively. Green: rotation axis. Black sphere: cell’s PD when the animal is upright. **‘Trajectory’:** head azimuth (in TA and YO frames: blue and cyan, respectively) and tilt (red) resulting from a rotation of ±180° relative to the 0° position. **’Tuning Curves’:** simulations of each model are compared with experimental results. Firing rate is represented in polar coordinate for consistency with (Shinder and Taube 2019) and normalized to 1 (green circle) when facing the PD in upright orientation. Blue: simulated firing of a cell encoding TA azimuth; Cyan: simulated firing rate of a cell encoding YO azimuth; Black: experimental results of Shinder and Taube (2019). When the trajectory corresponds to multiple manipulations, the corresponding results are averaged. **’YO vs TA’:** correlation between the tuning curves predicted by the YO and TA models and the average HDC response. **’Trajectory parameters’:** description of the trajectory, including head tilt, the position of the rotation axis in allocentric and egocentric coordinates, and decomposition of the rotation axis based on the dual-axis rule.

In the next two manipulations, the head it tilted 45° in pitch (**Fig. 4B**) or roll (**Fig. 4C**), such that the rotation axis now falls between the head’s yaw and roll axes (**Fig. 4B**) or yaw and pitch axes (**Fig. 4C**). According to the TA model, this rotation corresponds to the second updating rule (axis parallel to gravity), and the brain integrates 3D (yaw, pitch, roll) rotation signals to update azimuth accurately (‘Trajectory’ panels, blue). Because the head is tilted, HD tuning is reduced (**Fig. 3B**), although to a minimal extend since tilt angle is small (45°). Thus, the TA model predicts that cells exhibit a clear tuning (‘Tuning Curves’ panels, blue), in agreement with experimental results (ρ=0.98). In contrast, the YO model predicts that azimuth is not tracked accurately since the rotation is not aligned with the yaw axis. Specifically, yaw velocity is equal to 71% of the total velocity (i.e. the cosine of 45°) and therefore the YO model predicts that the rotation is underestimated: after a rotation of 360°, the estimated azimuth is 255° (‘Trajectory’ and ‘Tuning Curves’ panels, cyan), and therefore the simulated firing doesn’t return to the peak value which is expected when facing the PD. This prediction doesn’t correlate well (ρ=0.6) with the average tuning curves reported by Shinder and Taube (2019).

In the subsequent manipulations (**Fig. 4D,E**), head tilt increases to 90° such that the rotation occurs in the pitch or roll plane. As in the previous manipulations, the TA model predicts that these pitch and roll rotations correspond to the 2^nd^ updating rule and are thus integrated to generate a veridical azimuth signal (‘Trajectory’ panels, blue), although the cell’s response tuning is now significantly attenuated due to the large head tilt (‘Tuning Curves’ panels, blue). In contrast, the YO model predicts that HDC do not detect any change in azimuth (‘Trajectory’ panels, cyan), and therefore the simulated firing rate is identical at all head positions (‘Tuning Curves’ panels, cyan). When averaged across all manipulations, we found that HDC responses measured by Shinder and Taube exhibited a weak modulation that matched the TA model’s simulation (ρ=0.86/0.95 in **Fig. 4D/E**, respectively). In contrast, Shinder and Taube’s YO model predicts that HDC should not respond at all to these manipulations, in contradiction with their own results (in this case, the correlation is undefined). Remarkably, although these manipulations provide strong experimental support for the TA model, the authors reached the opposite conclusion without any simulations or quantification!

In a final manipulation (**Fig. 4F**), the head is placed upside down. In this simulation, both models predict that the neuron is unmodulated.

#### HDC responses during yaw rotations in an earth-vertical plane

Next we consider a series of manipulations where the animal rotates in yaw about an earth-horizontal axis. These manipulations cannot distinguish between the TA and YO models, because predictions are identical: both models predict that azimuth is updated by the rotation, although the predicted firing is attenuated since the head is tilted by 90° (**Fig. 5A-C**, ‘Trajectory’ panels).

**Figure 5:**
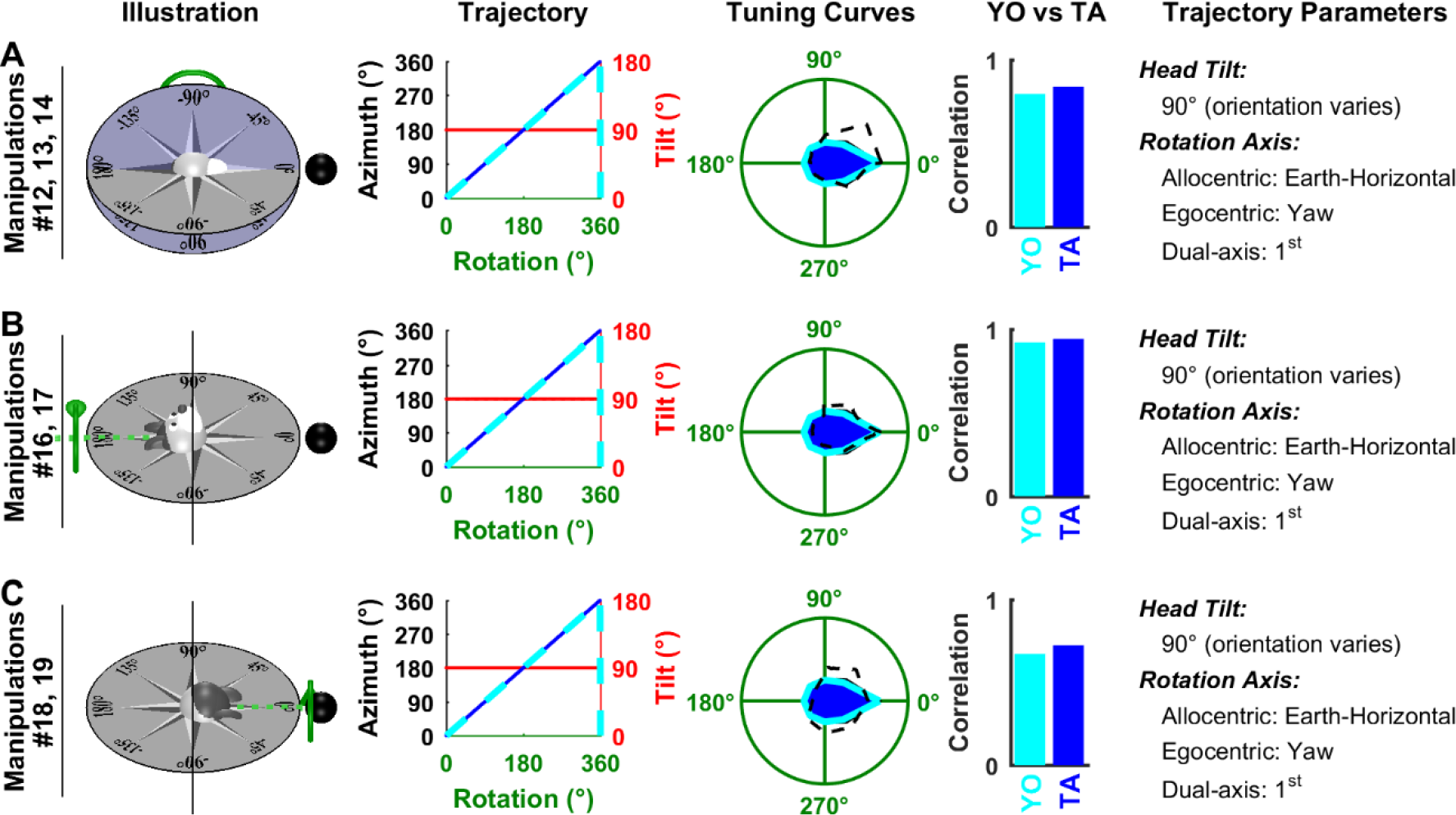
Analysis of Shinder and Taube’s (2019) manipulations: yaw rotations in the earth-vertical plane. **A**: the top of the head faces away from the reader. **B,C:** the top of the head faces left/right. In all panels, the head rotates in yaw. Same format as in **Fig. 4**.

The predicted tuning curves match equally well the experimental data (TA model: ρ=0.84/0.94/0.72; YO model: ρ=0.79/0.92/0.67).

#### HDC responses during pitch and roll rotations in an earth-vertical plane

Shinder and Taube also performed a series of manipulations where animals rotate in pitch or roll in the earth-vertical plane. According to the TA model, these rotations are about the null axis and should therefore not affect azimuth. According to the YO model, these rotations are about an axis orthogonal to the yaw axis and shouldn’t affect azimuth either. Thus, these manipulations don’t contribute to distinguishing these models. Yet, they provide additional support to our modeling framework, and in particular to the tilt-dependent gain function used here (**Fig. 3B**).

In **Fig. 6A,B**, the animal faces the cell’s PD at the beginning of rotation. Since azimuth doesn’t change during the whole rotation cycle, the animal faces the PD during the entire manipulation (‘Trajectory’ panels, blue and cyan curves have a constant value of 0). However, the models predict that the cell’s response is modulated by tilt angle (**Fig. 3B**), resulting in a broad tuning curve (‘Tuning curves’ panels, blue and cyan) that matches the average responses reported by Shinder and Taube (TA model: ρ=0.84 and ρ=0.76 in **Fig. 6A** and **B**; YO model: ρ=0.85 and ρ=0.76).

**Figure 6:**
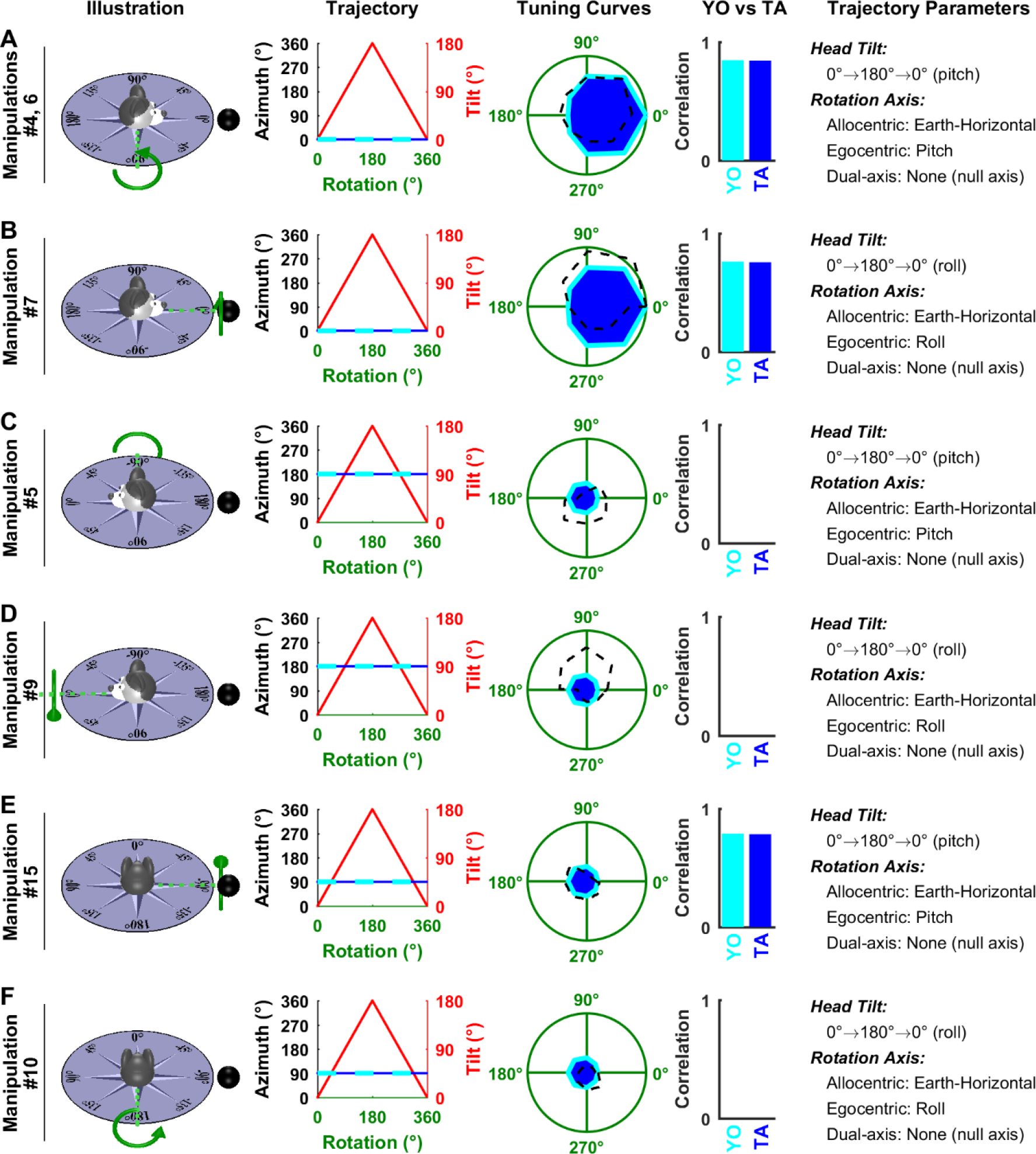
Analysis of Shinder and Taube’s (2019) manipulations: pitch/roll rotations in the earth-vertical plane. Same format as in **Fig. 4**. Note that, strictly speaking, TA (‘Trajectory’ panels, blue) is not defined when the tilt angle (red) is 180°. However, it is defined in the immediate vicinity of this point, such that the blue curve appears uninterrupted.

In **Fig. 6C-F**, the animal initially faces 90° or 180° away from the cell’s PD. As in previous manipulations, both models predict that the rotation doesn’t change azimuth (‘Trajectory’ panels, blue and cyan curves). The cell’s predicted response corresponds to its firing at 90° or 180° azimuth in **Fig. 3B** and is affected to a small extent by tilt angle. Shinder and Taube (2019) observed a weak modulation in **Fig. 6C,D**, that wasn’t predicted by either the TA or YO model (but might be attributable to tilt tuning, see next section). In **Fig. 6E,F**, the average firing was weak, as predicted by both models. In **Fig. 6E**, it correlated with the predicted tuning curve, although this correlation may not be very meaningful due to the weakness of the predicted modulation.

#### Other manipulations

Shinder and Taube (2019) also performed complex rotation protocols illustrated in **Fig. 7**. In **Fig. 7A**, the head starts from 45° nose-up tilt and rotates around an earth-horizontal axis. In this situation, TA follows a non-linear trajectory (‘Trajectory’ panel, blue) while the YO model predicts that rotation velocity is underestimated (‘Trajectory panel’, cyan), similar to **Fig. 4B,C**. The average measured firing rate measured correlates better with the TA than YO model (ρ=0.86 vs 0.52).

**Figure 7:**
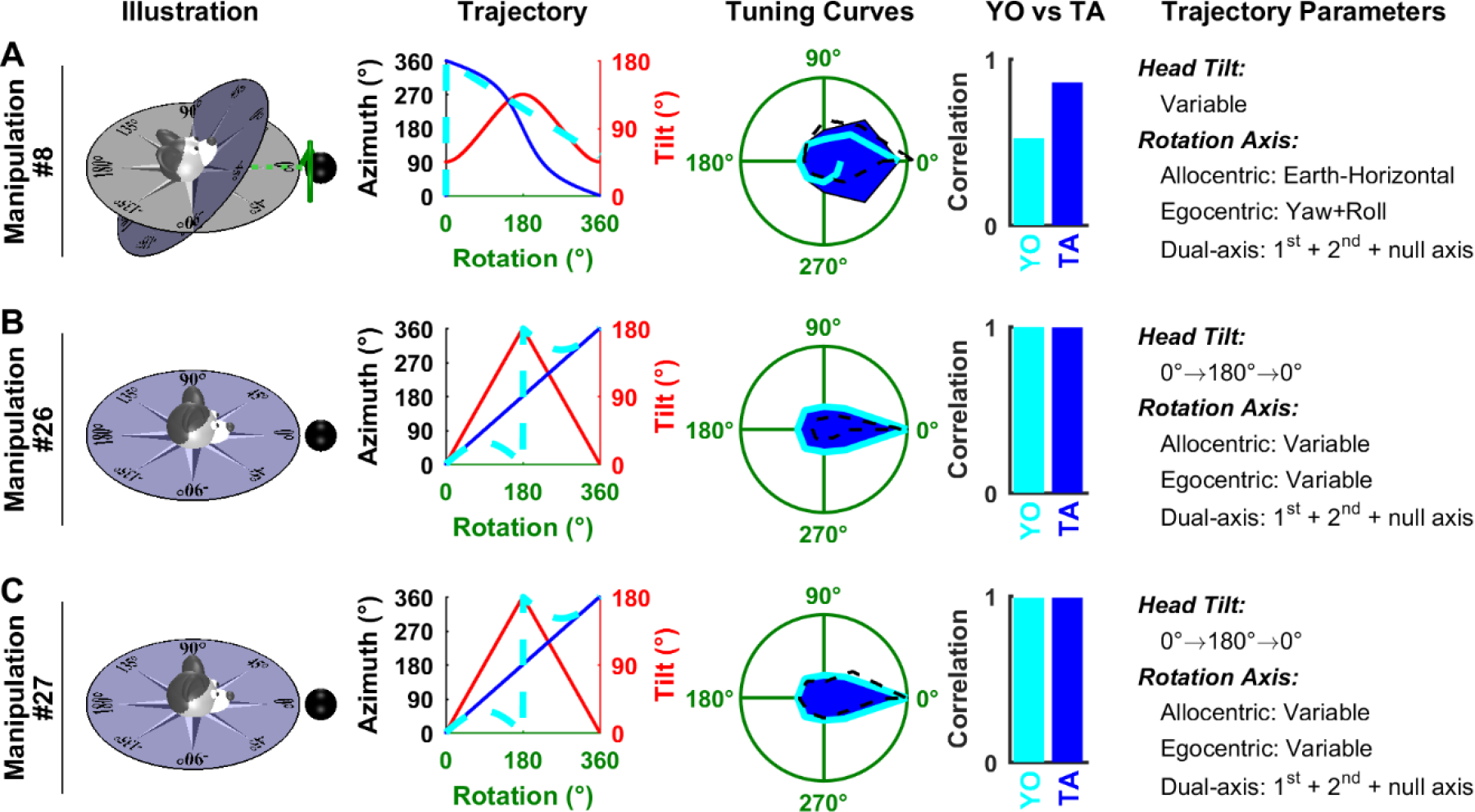
Analysis of Shinder and Taube’s (2019) manipulations: complex rotations. Same format as in **Fig. 4**. As in **Fig. 6**, TA is not defined at 180° tilt in panels B,C; but the corresponding curve (‘Trajectory’ panels, blue) appears uninterrupted since TA is defined in the vicinity of this point.

The manipulations in **Fig. 7B,C**, are combinations of yaw and roll or yaw and pitch (that can’t be represented using a single rotation axis as in other panels). The TA and YO models predict that azimuth follows different trajectories (‘Trajectory’ panels, blue versus cyan). However, in both trajectories, the head is inverted after 180° of rotation, resulting in a low firing rate, and the head returns to facing the PD after 360° of rotation, resulting in a high firing rate. Therefore, the predicted responses are very similar (‘Tuning curves’ panel, blue versus cyan), despite the trajectories being different. Both models predict that HDC should be strongly modulated during these simulations, in agreement with simulation results (TA model: ρ= 0.99 and 0.99 in **Fig. 7B** and **C**; YO model: ρ=1.00 and 0.99 in **Fig. 7B** and **C**).

### Conclusions: Shinder and Taube’s experimental results support the TA, and not the YO model

One of the two main conclusions of Shinder and Taube (2019) is that azimuth is only updated by yaw rotations, using incorrect qualitative arguments, rather than model-based analyses. In contrast, here we have simulated each of the two (TA and YO) models and compared with their experimental data. Strikingly, this comparison demonstrates that the experimental data is rather supportive of the TA/dual axis rule model, where HDC encode TA and, thus, by definition, integrate rotations about all three head axes. We may also point out that, while the attenuation of HDC tuning is a complicating factor, both the TA and YO models assume its existence. Thus, this factor doesn’t affect the comparison between the two models.

The critical manipulations for distinguishing between the YO versus TA models are rotations about an earth-vertical axis (which represents the 2^nd^ axis of the dual axis rule) with the animal at different static orientations, as shown in **Fig. 4**. Shinder and Taube’s YO model predicts that HDC should not be updated correctly in **Fig. 4B,C**, when the head is tilted by 45° and the rotation is misaligned with the yaw axis, and that tuning should vanish altogether in **Fig. 4D,E**, when the head is tilted by 90° and the rotation occurs along the pitch or roll axis. However, their data indicate that the average HD tuning is maintained in **Fig. 4B,C** as well as in **Fig. 4D,E** although it is attenuated in the later conditions because the head is tilted. In their study, Shinder and Taube don’t discuss the fact that HDC tuning is maintained when the head is tilted 45° (**Fig. 4B,C**), a fact that contradicts the model they promote. Furthermore, they conclude from the manipulation in **Fig. 4D,E**, that HDC aren’t updated during rotations in the head’s vertical planes, without appreciating that the response attenuation may be due to head tilt, and not of the fact that the head rotates in pitch and roll.

To explore this matter further, we note that another veridical comparison among TA and YO model predictions without the possible contamination of tilt attenuation is provided by considering the manipulations in **Fig. 4D,E** and **Fig. 5**, where head tilt is identical. Based on the TA model, each of these rotations spans the whole azimuth compass, and, since head tilt is identical, the attenuation of the cell’s response should be identical. In contrast, the YO model predicts that HD cells should be tuned in **Fig. 5** but not at all in **Fig. 4D,E**. We averaged the experimental tuning curves in **Fig. 4D,E** and **Fig. 5** and found that the average tuning curves are similar (**Fig. 8**) and highly correlated (ρ=0.9), thus confirming the TA/dual axis model.

**Figure 8:**
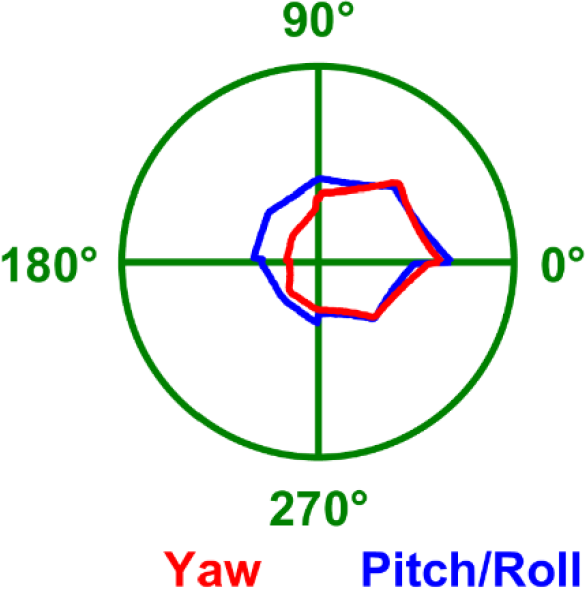
Average azimuth tuning curves measured by (Shinder and Taube 2019) with the head tilted 90°. Red: Tuning curve during yaw rotations in the earth-vertical plane (average of all manipulations in **Fig. 5**). This tuning is based on the 1^st^ axis rule. Blue: Tuning curve during pitch and roll rotations in the earth-horizontal plane (average of all manipulations in **Fig. 4D,E**). This tuning is based on the 2^nd^ axis rule. As predicted by the TA/dual axis model, the two curves are highly correlated (ρ=0.9). In contrast, the YO model would predict no tuning during pitch/roll (blue).

Finally, note that the tilt attenuation is directly responsible for the broad tuning curves measured in Fig. 6A,B, which are well fitted by both models. Thus, the existence of tilt attenuation, and our choice of a continuous gain modulation function to model it, are well supported by experimental data.

Shinder and Taube (2019) performed additional manipulations that resulted in elimination of directional tuning. For example, in conditions 12, 20, 24, 25, they observed that cells fired maximally at the beginning of the trajectory, when animals faced the PD, but that this firing didn’t recover after an entire 360° rotation when the animal returned to the PD. The authors emphasized that these occurred during rotations in pitch and roll (manipulations 20, 24 and 25), and actually used this point to support the YO model. However, they ignored the fact that a similar phenomenon also occurs in manipulation 12, which is a yaw rotation. Since loss of directional tuning may occur during rotations about all 3 head axes (yaw, pitch and roll), it doesn’t support any particular model (TA vs. YO). Instead, it likely reflects a decrease in the reproducibility of neuronal responses when the head tilts away from vertical. The absence of tuning in conditions 6 and 21 may be interpreted in a similar manner.

Shinder and Taube (2019) further reported that the average HDC responses may vary between different manipulations that follow the same trajectory but with different initial positions or directions (e.g. 12, 13, 14, **Fig. 5A**). However, this occurred only in trajectories where both models yielded identical predictions. Therefore, the variability of the responses across manipulations may involve neuronal response variability, alertness or other factors, but doesn’t weight in favor of a particular model.

Overall, although HD responses appear to be less consistent when the head tilts, responses that clearly support the TA frame and dual axis rule were observed at least in one manipulation for each trajectory. Furthermore, Shinder and Taube (2019) never observed a strong directional tuning in a direction that was not predicted by the TA model. Therefore, there is absolutely no evidence in their experimental results in favor of the YO model.

Note that, in our model analysis, we reproduced the average tuning curve, across all recorded cells, by simulating an HD cell that was tuned to azimuth, but not to tilt. This doesn’t imply that individual cells recorded by Shinder and Taube (2019) were not tuned to tilt, but rather than tilt tuning would generally average itself out when data from multiple cells, that would likely prefer different tilt, are pooled. We discuss tilt tuning in detail next.

### Tilt responses shown in (Shinder and Taube 2019) are biased by azimuth tuning

A second conclusion of Shinder and Taube (2019) is that HDC don’t encode head tilt in vertical planes. This conclusion is based on the analysis of pitch and roll responses recorded when the head initially faced the PD (Fig. 6 and 7 in their study; i.e. manipulations 4 and 7; **Fig. 6A,B** in the present manuscript). Specifically, the authors found that neuronal responses were systematically higher when the animals are upright, compared to responses at tilts larger than 90°, and concluded that HDC don’t exhibit any preference for any tilt position other than upright.

Here we show that this observation is biased because the cells’ azimuth tuning decreases as a function of tilt. To illustrate our reasoning, we first define and incorporate tilt tuning in our model, then simulate HDC responses for conjunctive tilt and azimuth tuning.

### Tilt tuning

Recall that we have defined tilt-dependent azimuth tuning curves such that the average value, across all azimuth angles, is constant (**Fig. 9A**, tilt marginal). Now we define tilt tuning as the cell’s firing rate as a function of tilt angle, averaged across all azimuth angles **(Fig. 9B-D)**. Thus, a cell whose firing rate is sampled uniformly and, when averaged across azimuth, is independent of tilt, would be classified as azimuth-tuned, but not tilt-tuned. In contrast, tilt tuning should manifest itself as an increase in the cell’s firing at a certain tilt angle, regardless of azimuth (**Fig. 9B**). To analyze Shinder and Taube’s results, we only need to simulate tilt tuning during rotations in a single vertical plane, i.e. pitch or roll. For simplicity, we model tuning curves along a single rotation axis (e.g. pitch) using a von Mises function:

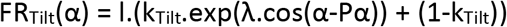

where λ is the coefficient of the von Mises distribution, k_Tilt_ a gain factor, α is the tilt angle, equal to 0 or 360° in upright position and 180° when upside-down. The parameter l is a scaling factor adjusted so that the curve’s peak value is 1.

**Figure 9:**
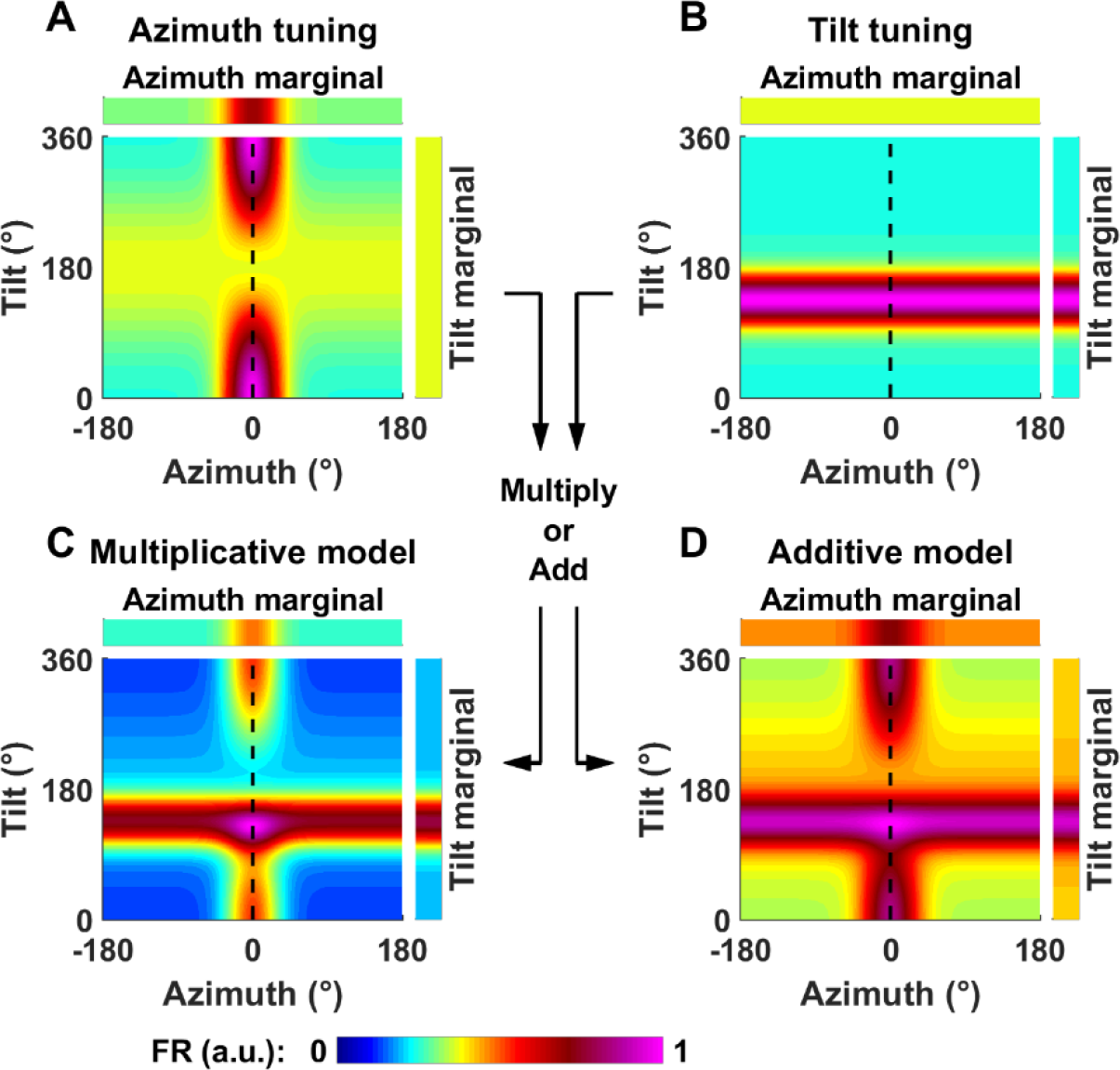
Definition of tilt tuning and generalization to conjunctive azimuth and tilt tuning. **A: Model cell tuned to azimuth only**. Tilt-dependent azimuth tuning curve of a model cell tuned to azimuth only (as in **Fig. 3A**, with the tilt axis ranging from 0° to 360°). **B: Model cell tuned to tilt only**. Tilt tuning is independent of azimuth and peaks at 135°. **C-D: Model cell tuned conjunctively to both azimuth and tilt**. Conjunctive tuning curve, assuming that tilt and azimuth tuning interact either multiplicatively (C) or additively (D). These simulations assume tilt and azimuth tuning of equal strength. That is, simulation parameters of tilt tuning curve (λ = 4.6, k_Tilt_ = 0.46) are identical to the parameters κ and k_G_ of the azimuth tuning curve. Note that, for simplicity, we express tilt along one axis only here (e.g. pitch or roll) and assume that the cell responds preferentially at a certain tilt angle (135°) along this axis.

To simulate the response of a cell that encodes both azimuth and tilt conjunctively, we consider two alternatives where tilt and azimuth interact multiplicatively or additively:

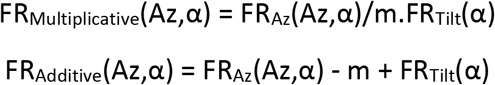

In both equations, m is equal to the average of FR_Az_(Az,α) across all azimuth angles (which is independent of α). It is introduced in both equations to ensure that the average values of FR_Multiplicative_(Az,α) and FR_Additive_(Az,α) across all azimuths are equal to FR_Tilt_(α), in agreement with the definition of the tilt tuning curve.

Example conjunctive tuning curves, assuming multiplicative or additive interaction, are shown in **Fig. 9C,D**. In these examples, we have assumed that azimuth and tilt tuning have identical strength. In this case, the resulting 2D tuning curve adopts a ‘cross’ shape, with a horizontal band corresponding to an increased firing at the preferred tilt angle and a vertical band corresponding to an increased firing at the preferred azimuth.

### Bias by azimuth tuning

To illustrate how azimuth tuning may bias tilt tuning identification, we simulate another conjunctive cell where azimuth tuning (simulated with κ = 2 and k_G_ = 1; **Fig. 10A**) is stronger than tilt tuning (simulated with λ = 0.5 and k_Tilt_=1; **Fig. 10B**). We assume that tilt tuning peaks at 135° (**Fig. 10B**, white line).

**Figure 10:**
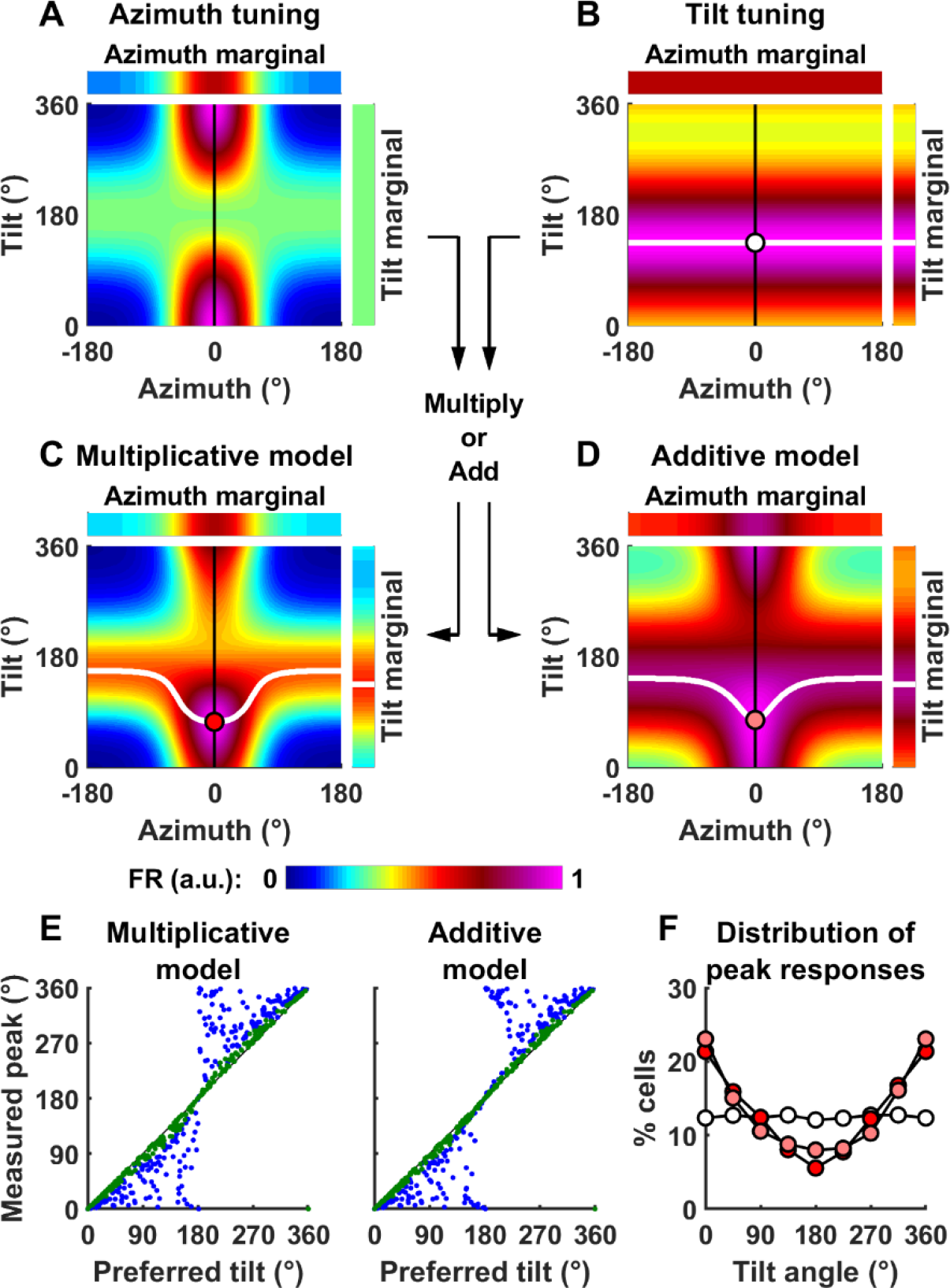
Why the pitch/roll rotations when facing the cell’s PD (used by Shinder and Taube) are inappropriate to test for tilt tuning. **A-D:** Simulation of a model conjunctive cell, as in **Fig. 9**, but with different parameters such that azimuth tuning (κ = 2 and kG = 1) is now stronger than tilt tuning (λ = 0.5 and k_Tilt_ = 1). The solid white lines in B-D indicate the tilt angle at which firing peaks, as a function of azimuth. The tilt tuning function in B, where tilt tuning is independent of azimuth and peaks at 135°, is multiplied by, or added to the azimuth tuning function (A) to produce the tuning curves in C and D. The white (B), red (C) or pink (D) markers indicate the tilt angle at which firing peaks when facing 0° azimuth, i.e. the cell’s PD: note the bias towards upright. **E:** Peak response measured during pitch or roll, as a function of the cell’s preferred tilt angle, in 500 simulated cells (k_G_ and are k_Tilt_ set to 1, κ and λ are drawn randomly such that the Rayleigh vector length of azimuth and tilt tuning curves are distributed uniformly). Green/blue dots: cells where tilt tuning is higher/lower than azimuth tuning. **F:** Distribution of the preferred tilt angle (white markers) and of the angle at which the peak response occurs (red/pink: multiplicative/additive model).

Under these conditions, azimuth and tilt tuning interact to form a response peak at 0° azimuth and, importantly, at a smaller tilt angle than 135° (specifically at a tilt angle of α=74° and α=77° in **Fig. 10C,D** red/pink circle respectively). This is because the tilt-dependent azimuth tuning (**Fig. 10A**), which is maximal at 0° azimuth and near upright, leads to higher firing rates for small tilt angles. Thus, the fact that azimuth tuning strength is tilt-dependent results in biases of response peak towards upright, even though tilt PD is at 135°. Note that this ‘pulling’ of response peak towards upright is maximal for tilts along the azimuth PD (0° in the present simulations) (**Fig. 10C,D**, white lines) – and these are exactly the conditions tested by Shinder and Taube (2019).

The extent to which azimuth tuning biases the response to tilt tuning depends on the relative strength of azimuth vs. tilt tuning (Rayleigh vector length of the corresponding marginals: 0.79 versus 0.25 in **Fig. 10A,B**). We modeled a population of neurons where the tuning strength of tilt and azimuth (measured by their Rayleigh vector length) are independent and distributed uniformly between 0 and 1, and where the preferred tilt is distributed uniformly between 0 and 360° (**Fig. 10E**, abscissae). We repeated the simulations in **Fig. 10C,D** and computed the tilt angle at which firing rate would be maximum (**Fig. 10E**, ordinate axis), when facing 0° azimuth, and assuming a multiplicative (left) or additive (right) interaction. Furthermore, we color-coded neurons where tilt tuning was stronger than azimuth tuning in green. These neurons appear close to the diagonal, indicating that pitch tuning was weakly biased in these neurons. In contrast, the peak response of neurons where azimuth tuning is larger than tilt tuning (blue) appeared away from the diagonal and close to 0 or 360° on the ordinate axis, indicating that it is strongly biased towards upright. Note that Shinder and Taube only recorded from cells with strong azimuth tuning, thus inherently biasing their data in the direction of the blue symbols.

To summarize the population responses, we plotted the distribution of preferred pitch orientation in the absence of azimuth tuning (**Fig. 10F**, white symbols) and the distribution of peak firing during pitch rotation assuming a multiplicative (red) or additive (pink) model. The later distributions are biased towards 0°, i.e. upright.

Furthermore, additional sources of bias should be considered:

- First, Shinder and Taube recorded only azimuth-tuned HDC with a Rayleigh vector length higher than 0.5. Therefore, cells were pre-selected to have a high azimuth response, but were not selected to have a large tilt response, implying that most cells would have a stronger azimuth tuning than tilt tuning (i.e., blue symbols in **Fig. 10E**). Since HDC with a larger azimuth tuning compared to tilt tuning are biased towards responding maximally in upright, this pre-selection would have biased their results further.
- We simulated rotations in a single plane (e.g. pitch) and assumed that the tuning strength of tilt tuning is uniformly distributed, similar to azimuth tuning. However, tilt-tuned cells may in fact respond preferentially in a different plane, e.g. roll. Because of this, the distribution of tilt tuning strength should be biased towards lower values and the population response should be biased even further towards upright.

Given these multiple sources of bias, and considering the limited set of cells analyzed (only 11 and 13 cells in manipulations 4 (pitch rotation) and 7 (roll rotations), respectively), it is not surprising that Shinder and Taube didn’t observe any cells firing maximally in the range of 135-215° tilt, i.e. within ±45° of being upside-down (although some of their cells fired maximally at 90° or 270° tilt, i.e. half-way between upright and upside-down).

### Conclusion: Tilt tuning

We conclude that the manipulations shown and analyzed in Shinder and Taube (2019) are insufficient to characterize the presence or absence of tilt tuning. Our simulations indicate that they would have observed a clear tilt tuning (characterized by an increase in firing rate at tilt angles larger than 90°), only in cells where (1) tilt tuning is stronger compared to azimuth tuning, which would likely represent a minority of the population since recordings were performed in cells where azimuth tuning was strong in the first place, (2) the preferred tilt angle is larger than 90°, and (3) tilt tuning occurs in the plane in which recordings were performed. Considering the limited sample shown in Shinder and Taube (Figures 6 and 7 in their study: 11 and 13 cells respectively), it is no surprising that no such cells were found.

Tilt responses would be better characterized by measuring pitch and roll responses when the head faces 90° or 180° away from the preferred azimuth (**Fig. 6C-F** in the present manuscript). Although this data was collected by Shinder and Taube (2019), it was not used to assess the presence or absence of tilt tuning and individual cell responses were not shown.

## Discussion

### Shinder and Taube have drawn two conclusions from their dataset

The first conclusion is that the azimuth tuning of HDC follows a ‘yaw-only’ model (**Fig. 1A**), as stated in their discussion: *“Because the system could not effectively utilize non-horizontal information to determine rotation in the horizontal plane, this result challenges the viability of the internal model, which postulates that the brain uses all available sensory data in combination with gravity information to derive a directional heading vector. […] Instead, our findings suggest that the horizontal canals and its associated pathways are hard-wired and designed to specifically extract azimuthal heading information – most likely in the form of angular head velocity information.”*. We have shown that this conclusion is not supported by their data. In fact, a model-based analysis of their experimental protocols and neural responses shows the exact opposite: HDC responses are inconsistent with the YO model, but instead support the TA/dual axis model (Page et al. 2017; Laurens and Angelaki 2018), where 3D (yaw, pitch, roll) rotation information originating from all semicircular canals are integrated with gravity signals to track azimuthal heading. In particular, we showed that pitch and roll rotations are not expected to update azimuth based on either the TA or YO model in most manipulations performed by Shinder and Taube (**Fig. 6**). However, in those manipulations where they should update azimuth according to the dual axis rule (e.g. **Fig. 4B-E**), they do it just as effectively as yaw rotations (see **Fig. 8**).

The second conclusion is that HDC don’t encode head tilt, as stated in their discussion: *“HD cells increased their firing rates when the animal faced into the recorded cell’s PFD in the horizontal plane, independent of the tilt or roll position of the head, as long as the head did not become inverted by tilt or roll beyond 90°”*. We agree with their conclusion that azimuth signals carried by HDC decrease with head tilt (although likely not abruptly, as implied by the hemitorus and ellipsoid but never supported by any data) but we showed that their data is inconclusive regarding whether HDC are tuned to tilt.

The YO model proposed by Shinder and Taube implies that HDC encode exclusively 1D information. According to the authors, the only effect of 3D motion on HDC is that azimuth tuning vanishes when animals are inverted. The authors do not describe this as a coding strategy but merely as an indication that otolith signals play a role in computing head orientation, and clearly state that HDC do not, in their view, encode pitch or roll tilt: *“whereas a sizeable percentage of HD cells in bats (30%) were found to have conjunctive properties with pitch and roll, where the cells were best tuned to a particular orientation in 3D space and not just in the azimuthal plane (Finkelstein et al., 2015, Fig. 4G), there was limited evidence to support this same occurrence in the rat anterodorsal thalamus”*. Therefore, we find it surprising that the authors would state, in their study’s abstract and title, that “the HD signal is a 3D gravity-referenced signal”, in what seems in blatant contradiction with their own conclusions.

There are other misleading – in fact, erroneous – statements; e.g., “*In this regard [referring to the fact that directional firing is disrupted during inversion], the hemitorus and ellipsoid models when considered fixed to gravity are similar to the dual-axis model (Page et al. 2018), which postulates that the reference frame for HD cells is defined by two parameters: the animal’s head position relative to its dorsal-ventral axis and the relationship of the animal’s dorsal-ventral axis to gravity*”: this implies that the YO and TA models are identical because both somehow (and loosely) depends on gravity. This is bluntly incorrect; as illustrated here, the two models are entirely different. This statement also misses the crucial point that the dual-axis model requires integration of 3D rotation signals, thus contradicting their own conclusion that HD cells integrate yaw rotations only. The YO and TA/dual axis models are identical <u>only</u> when upright or moving in 2D (i.e., walking up and down vertical walls, without transitioning between different vertical walls; **Fig. 1A**). They differ entirely in every other respect: both in their functional significance (e.g., the YO model loses allocentric invariance during 3D re-orientations) and in the type of multisensory signals that are necessary to define and update HDC tuning.

We have proposed (Laurens and Angelaki 2018) that the HD system combines visual landmark signals with self-motion information provided by a multisensory 3D internal model (Merfeld et al. 1999; Laurens and Droulez 2007; Laurens and Angelaki 2011, 2017; Laurens et al. 2013). The fact that HDC responded as predicted (**Fig. 4-7**) along all three rotation axes supports the 3D internal model theory. Note that all manipulations in the Shinder and Taube (2019) study were performed in light, and likely involve a combination of vestibular self-motion signals, visual self-motion signals, and visual landmarks. Therefore, HDC responses didn’t reflect how the brain processes vestibular signals specifically, but rather how it maintains a 3D representation of head orientation based on multisensory signals.

In nearly two decades (Stackman et al. 2000, Calton and Taube 2005, Shinder and Taube 2019), the conclusions of Shinder and Taube’s group, which is that HDC integrate yaw rotations exclusively to encode 1D azimuth, have remained unchanged. Several groups (Page et al. 2017, Laurens et al. 2016, Angelaki et al. 2016; Cham et al. 2017; Laurens et al. 2017; Finkelstein et al. 2015) have recently discovered that HDC encode 3D head orientation and derive orientation signals from 3D rotation signals. Here we have shown that the Shinder and Taube (2019) experimental results are entirely consistent with the conclusions of these other studies, although the authors failed to assimilate recent developments to interpret their data in a more model-driven, quantitative way.

## Acknowledgements

This work was supported by Simons Foundation Grant 542949 and Simons Collaboration on the Global Brain R01-AT010459

